# Entomophthovirus: An insect-derived iflavirus that infects a behavior manipulating fungal pathogen of dipterans

**DOI:** 10.1101/371526

**Authors:** Maxwell C. Coyle, Carolyn N. Elya, Michael Bronski, Michael B. Eisen

## Abstract

We discovered a virus infecting *Entomophthora muscae*, a behavior-manipulating fungal pathogen of dipterans. The virus, which we name Entomophthovirus, is a capsid-forming, positive-strand RNA virus in the viral family iflaviridae, whose known members almost exclusively infect insects. We show that the virus RNA is expressed at high levels in fungal cells *in vitro* and during *in vivo* infections of *Drosophila melanogaster*, and that virus particles are present in *E. muscae*. Two close relatives of the virus had been previously described as insect viruses based on the presence of viral genomes in transcriptomes assembled from RNA extracted from wild dipterans. By analyzing sequencing data from these earlier reports, we show that both dipteran samples were co-infected with *E. muscae*. We also find the virus in RNA sequencing data from samples of two other species of dipterans, *Musca domestica* and *Delia radicum*, known to be infected with *E. muscae*. These data establish that Entomophthovirus is widely, and seemingly obligately, associated with *E. muscae*. As other members of the iflaviridae cause behavioral changes in insects, we speculate on the possibility that Entomophthovirus plays a role in *E. muscae* involved host manipulation.

## Introduction

A wide variety of microbes have evolved the ability to manipulate animal behavior in ways that appear to advance microbial fitness (Eisthen and Theis, 2016; Forsythe et al., 2012; Hoover et al., 2011; Hughes et al., 2016; Libersat, 2003; Rohrscheib and Brownlie, 2013; Roy et al., 2006; Sampson and Mazmanian, 2015; Wang et al., 2015). Among them is the fungal pathogen of dipterans *Entomophthora muscae*. Originally identified in the 19th century (Cohn, 1855), *E. muscae* has been observed infecting a wide variety of fly species (Kramer and Steinkraus, 1981; Steinkraus and Kramer, 1987), which exhibit a distinct series of behaviors prior to death: they climb to a high location (summiting), extend their proboscides which become attached to the surface via fungal growths that protrude from the tip (Balazy, 1984), and extend their wings in a characteristic “death pose.”

Shortly after flies killed by *E. muscae* die, conidiophores emerge through the weakest points of the cuticle and forcibly eject spores at high velocity, ideally (from the fungal point of view) landing on a new host and propagating the infection. The wing position removes a major obstacle to spores escaping the immediate vicinity of the fly, and the elevation benefits the fungus by increasing the target range covered by traveling spores.

We recently reported the isolation of a strain of *E. muscae* from wild *Drosophila* and its propagation in lab-reared *D. melanogaster* and as an *in vitro* culture (Elya et al., 2017). *Drosophila* infected by this strain of *E. muscae* manifest the same set of behavioral changes as have been described in other flies.

In order to build a reference *E. muscae* transcriptome free of *Drosophila* RNA to study the behavior of the fungus in infected flies, we sequenced mRNA from our *in vitro* culture. As part of our initial quality checks of the *in vitro* mRNA sequence data, we used BLAST to search a small random subset of reads against GenBank for related sequences. We were surprised to find that a large number of reads from the *E. muscae* transcriptome aligned with near 100% identity to a virus identified by mRNA sequencing of wild *Drosophila* (Twyford virus; GenBank: [KP714075.1](https://www.ncbi.nlm.nih.gov/nuccore/821369846)) (Webster et al., 2015).

We initially suspected that this virus was misannotated - that the sample of flies from which the virus was isolated was infected with *E. muscae* and that the sequence was actually a transposable element from the repeat rich *E. muscae* genome. However when we subsequently began examining reads and initial assemblies of the *E. muscae* genome generated from our *in vitro* culture, we were astonished to find that the virus was not in our assembly or in any of the genomic reads from either Illumina or Pacific Biosciences sequencing.

This led us to more closely examine the original discovery of Twyford virus by (Webster et al., 2015). Twyford is one of roughly two dozen new viruses identified by assembling mRNAs isolated from multiple large collections of wild caught *Drosophila melanogaster* and screening for virus-like sequences not found in the *D. melanogaster* genome. It is an Iflavirus, a family of positive-strand RNA viruses, related to picornaviruses, that infect a wide variety of insect hosts. A nearly complete 8 kb genome of Twyford was assembled from flies caught in the southern England village that gave it its name.

Of the viruses isolated in this study two things stand out about Twyford. First, it is rare, only appearing once in a panel of 16 independent wild populations screened for known and newly identified *Drosophila* viruses. Second, and more notably, small RNAs derived from Twyford virus have unusual characteristics compared to those derived from other viruses: they show a strong negative strand bias and have an almost complete bias for a 5’ U base. Since this pattern was unique for *Drosophila* viruses, it suggested that small RNAs aligning to Twyford virus had not been generated by the canonical *Drosophila* Ago2-Dcr system. Webster et al. explored the hypothesis that the virus was infecting a eukaryotic commensal of *Drosophila*, but rejected candidate mites, nematodes and fungi for various reasons, and suggested the small RNAs they observed may have come from a previously unknown *Drosophila* pathway.

Here we present direct evidence that the virus they identified is infecting *E. muscae* and appears to be obligately associated with *E. muscae* in the wild. The viral RNA is found at high concentration in our *E. muscae* liquid culture, it cannot be washed away from fungal cells, and its expression tracks with *E. muscae* levels during infection of *Drosophila* in the lab. Small RNAs from our *in vitro* culture have precisely those characteristics described by ((Webster et al., 2015).

We purified the virus from our *in vitro* culture and have used electron microscopy (EM) to show that it forms structures of approximately the same size as other picornaviruses. We also used EM to show that there are intact virus particles in *E. muscae* cells, confirming that the virus is infecting the fungus.

We obtained reads from the original Twyford sample from public databases, and using our genomic data as a reference against which to screen reads, we show that at least one fly in that sample was infected with *E. muscae*. We also find close relatives of the virus present in three other transcriptomes of wild dipterans, two from individual flies known to be infected with *E. muscae*, and a third from a mixed collection of flies that also contains mRNAs from *E. muscae*.

Below we present details of our discovery, isolation and characterization of this virus, which we propose renaming Entomophthovirus to reflect its host and its novelty as the first picornavirus known to infect a fungus and the first virus of the insect-infecting iflaviridae known to infect a fungal pathogen of insects. We also discuss the possibility that the virus may be involved in behavior manipulation.

## Results

### Discovery of an iflavirus in in vitro culture of an isolate of *Entomophthora muscae*

We recently described the isolation of a strain of *E. muscae* from wild *Drosophila* species caught in Berkeley, CA in the summer of 2016. Following standard protocols (Hajek et al., 2012), we captured spores ejected from an individual *D. melanogaster* recently killed by *E. muscae* on a liquid medium (Elya et al., 2017). We verified that the culture contained *E. muscae* by genotyping both the ITS and 28S loci. We sequenced mRNA from the liquid culture approximately five months after it was established, obtaining 38.9 million paired-end reads of 150 bp.

As discussed above we ran individual reads from mRNAs from the *in vitro* culture through NCBI’s blastn, which we did to confirm that the sample was from *E. muscae*, and noticed that many reads aligned with ~95% identity to Twyford virus (GenBank: [KP714075.1](https://www.ncbi.nlm.nih.gov/nuccore/821369846)). After assembling an *E. muscae in vitro* transcriptome using TRINITY (Grabherr et al., 2011), we compared all of the assembled transcripts against Twyford, and found several highly similar versions of a slightly longer sequence (it appears the original Twyford virus sequence was truncated).

The genome of the virus in our liquid culture, which we refer to here as *D. melanogaster* Entomophthovirus (DmEV), is 8,832 basepairs and encodes a single 2,901 amino acid open reading frame. This single viral pro-protein contains the six proteins characteristic of iflaviruses: three coat proteins, an RNA helicase, a protease and a RNA-dependent RNA polymerase (Figure 1A). We built a phylogenetic tree to compare DmEV to other iflaviruses and picornaviruses, which shows its placement in a subclade with other insect viruses as well as a plant iflavirus (Tomato matilda virus) (Figure 1B).

**Figure 1:**
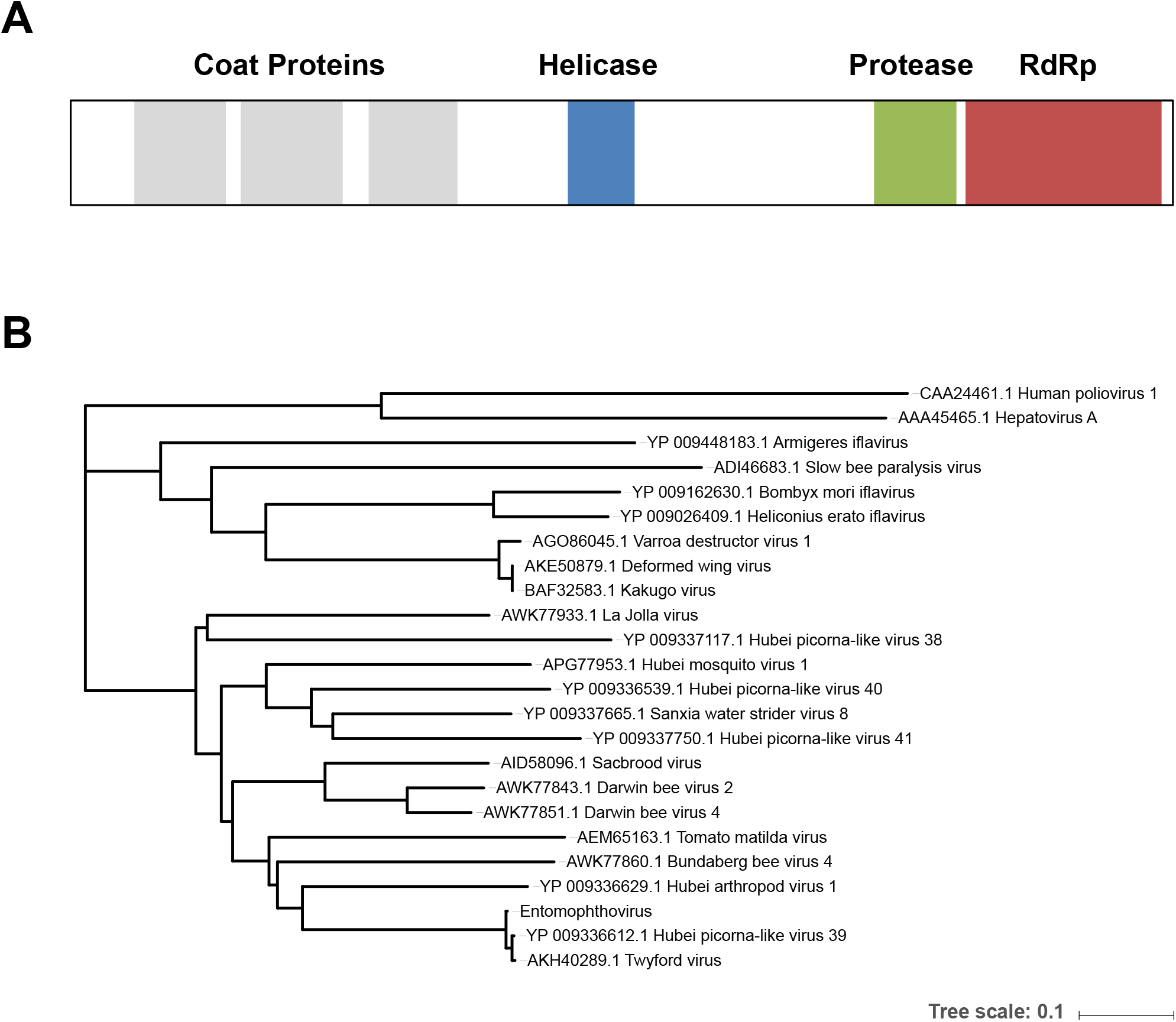
The genome and phylogenetic placement of Entomophthovirus. (A) The genome, assembled by Trinity from RNA-sequencing of our *in vitro Entomophthora muscae* culture, encodes a polyprotein with the characteristic open reading frames of a picornavirus, including three coat proteins, a helicase, a protease, and an RNA-dependent RNA polymerase. (B) A neighbor-joining phylogenetic re-construction of RdRp protein sequences clusters Entomophthorvirus with Twyford virus and Hubei picorna-like virus 39 as a sub-clade of the iflaviruses. Other characterized iflaviruses, including some linked to behavioral manipulation, are shown. The tree was calculated using MEGA7 (Kumar et al., 2016) using the Neighbor-Joining method (Saitou and Nei, 1987). The evolutionary distances were computed using the Poisson correction method (Zuckerkandl and Pauling, 1965) and are in the units of the number of substitutions per site. The optimal tree is shown, drawn to scale using iTOL (Letunic and Bork, 2016).

Reads aligning to this DmEV genome account for ~15% of the reads from the *in vitro* culture, suggesting that the virus is abundant and actively replicating within *E. musace* cells *in vitro*.

### Entomophthovirus expression during *in vivo* infection of *D. melanogaster* with *E. muscae*

We previously described an experiment in which we sequenced RNAs from either whole animals or extracted brains from *D. melanogaster* infected with *E. muscae* over the course of an infection, as well as time-matched uninfected controls (Elya et al., 2017). We examined reads from this experiment for DmEV and found significant levels of DmEV RNA in some flies infected with *E. muscae* (but not unexposed controls) beginning 72 hours post exposure (Figure 2), which is when we begin to see a significant fraction of reads aligning to the *E. muscae* transcriptome. In six of the twelve flies sampled 72 to 96 hours after infection, DmEV represents over ten percent of all reads (with a maximum of an astonishing 38 percent). There is, however, considerable inter-animal variation: at 72 hours between two and 21 percent of all reads align to DmEV, and at 96 hours the range is one to 38 percent. At 120 hours the fraction of reads aligning to DmEV drops while those aligning to *E. muscae* continues to rise.

**Figure 2.**
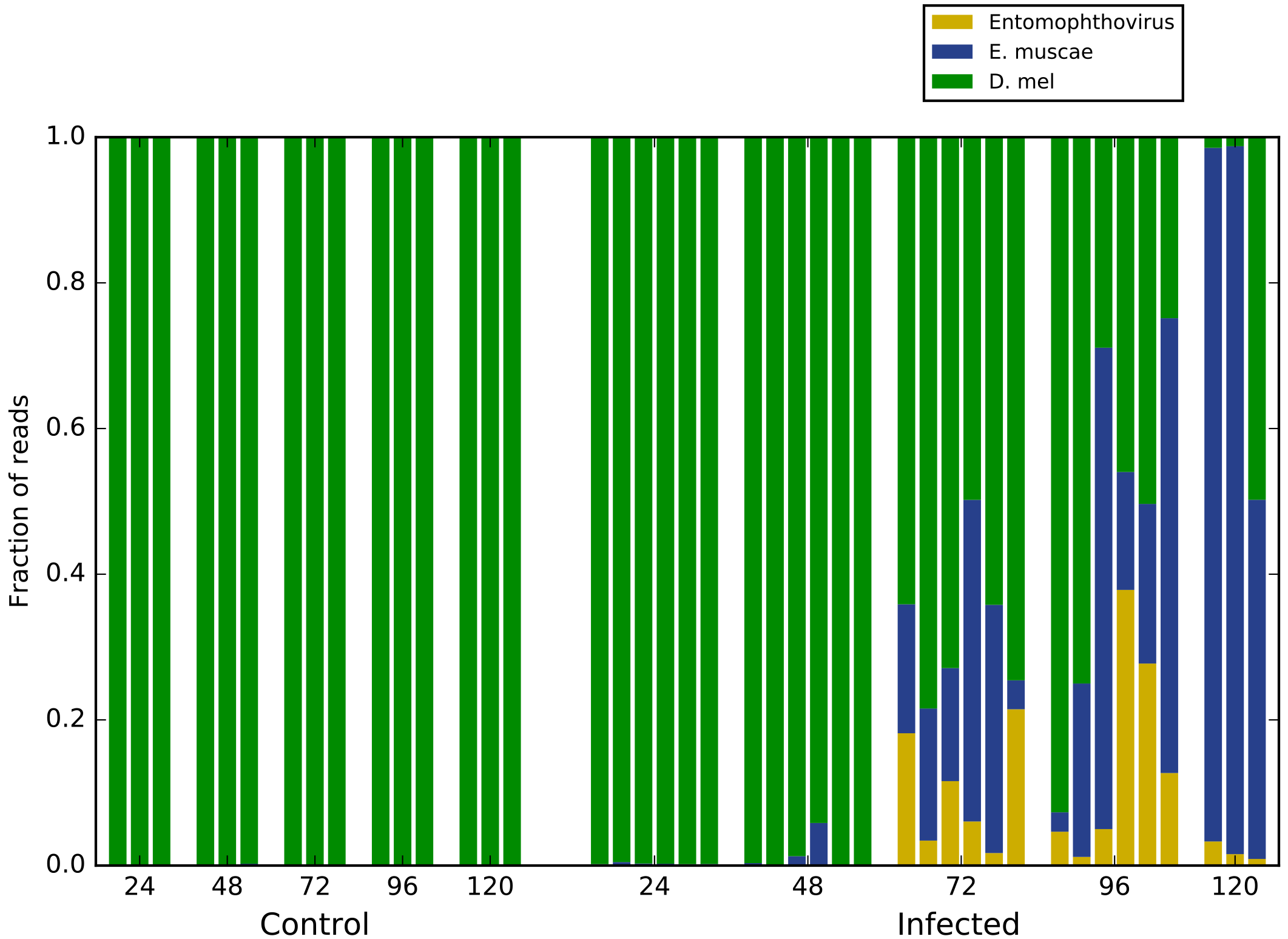
Entomophthovirus in *in vivo Entomophthora muscae* infections of *Drosophila melanogaster*. To characterize the growth of virus in animals infected with *E. muscae* aligned reads from mRNA-seq data of (Elya et al., 2017) to Entomophthovirus. Samples were from individual whole flies exposed to *E. muscae* at 24, 28, 72, 96 and 120 hours after infection as well as controls. Plotted are the fraction of total reads that aligned to *D. melanogaster* mRNAs, to *E. muscae* transcripts or annotated genes, and to Entomophthovirus.

### Evidence of intracellular and extracellular virus in *E. muscae in vitro* culture

At this point our only evidence for the existence of DmEV in the fungus was the presence of iflaviral RNA in a *Drosophila*-free *in vitro* culture, and we therefore sought to demonstrate that there are viral particles in the *in vitro* culture. We started with the supernatant, assuming that if viral particles are being made they would either actively exit cells or be released as cells are lysed.

We spun down fungal cells and filtered the supernatant through a 0.22 um filter to retain potential viral particles. Using primers to specifically amplify a 831 bp segment of the DmEV genome, we found DmEV enriched in the retentate by RT-PCR (Figure 3A). We then re-suspended the cell pellet and collected cells on a filter by vacuum filtration. After thoroughly washing the cells by vacuum filtration, we found a strong signal for DmEV in the eluted cell fraction (Figure 3A). No viral signal was detected in the media used to culture *E. muscae* or in stocks of the flies we use for our *in vivo E. muscae* infections.

**Figure 3.**
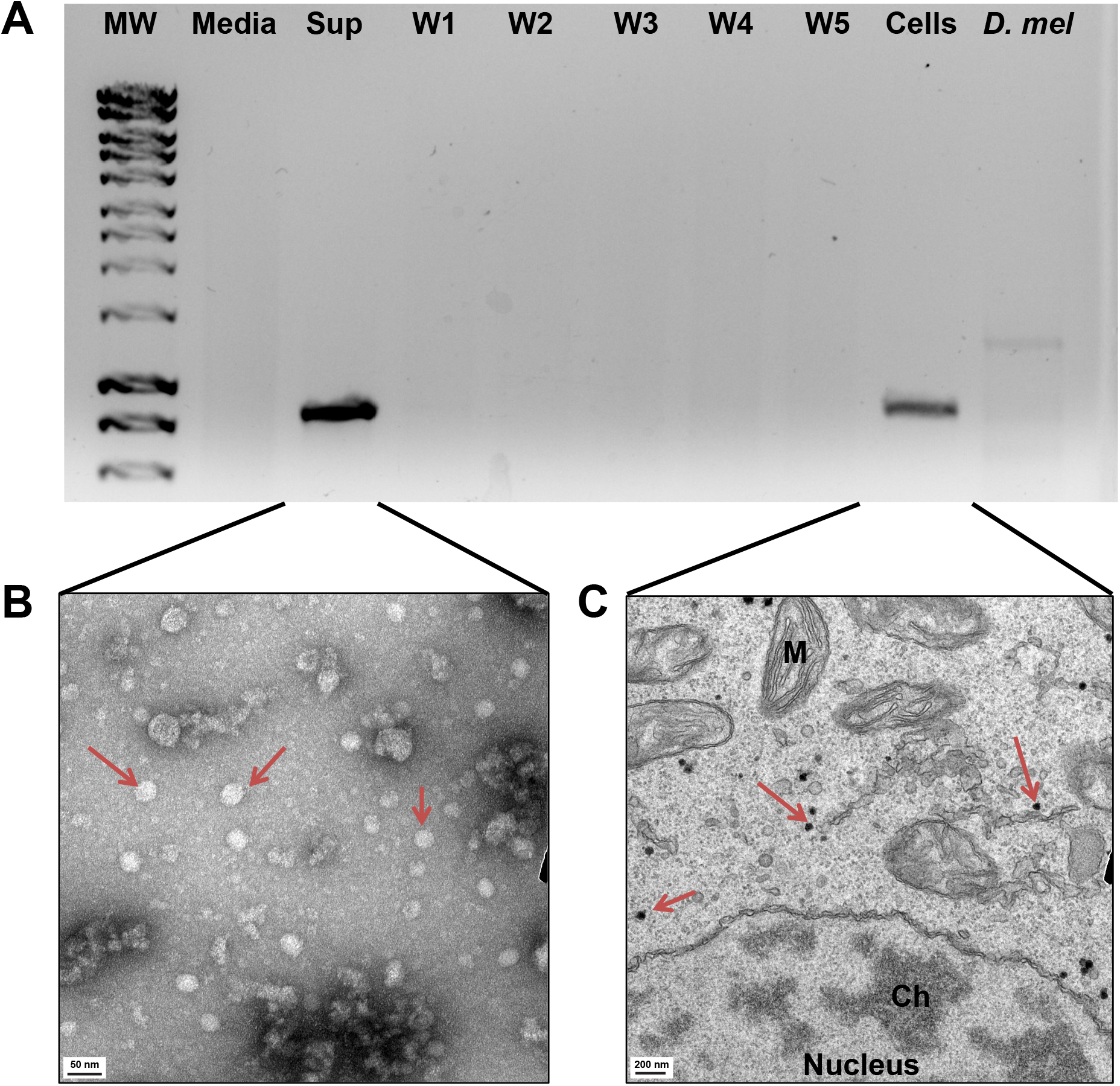
The presence of Entomophthovirus in *in vitro Entomophthora muscae* culture. (A) RT-PCR with primers specific for the Entomophthovirus genome shows the virus in the supernatant and cellular fraction of in vitro culture of *E. muscae*. Viral signal is lacking from the media we use for *in vitro* culture, the *Drosophila* stocks used to propagate the infection in lab, and washes of the cellular fraction. (B) Transmission electron microscopy of virus collected from the supernatant of in vitro cultures by ultra-centrifugation and negative stained by uranyl acetate. The viral particles (red arrows) have a tight size distribution and the expected diameter (~30 nm) of an iflavirus capsid. (C) Transmission electron microscopy of cellular sections of E. muscae, with a double-contrast staining of uranyl acetate and lead citrate. Electron-dense particles (red arrows) consistent with the size distribution of an iflavirus capsid are abundant in a fraction of cellular sections and are localized cytoplasmically. Examples of *E. muscae* nucleus, mitochondria (M), and chromatin (Ch) are also marked.

We used TEM to directly confirm the presence of virus particles, first in a sample purified from the *in vitro* supernatant (Figure 3B). Iflavirus capsids have been reported to have a diameter of around 30 nanometers (Silva et al., 2015), and the regular size and shape of viral particles help them stand out by TEM. Indeed, a uranyl acetate negative stain of viral particles collected by ultracentrifugation from the extracellular fraction showed an abundance of symmetric ~30 nm objects, consistent with our expectations for the iflaviridae (Fig 3B).

We next carried out double contrast uranyl acetate/lead citrate staining of fixed *E. muscae* cells to look for virus particles. A large fraction of sections had intracellular particles consistent with viral capsids by virtue of their ~30 nm diameter and their strong electron density (Figure 3C). Notably, these particles were never found inside the nucleus, mitochondria, or other clearly demarcated organelles (Fig 3C). Nor was their concentration noticeably higher near the plasma membrane or endomembranes, although the fixation process might have obscured fine spatial information. Only a fraction of sections seemed to possess any viral-like particles at all, while the infected cells showed a high viral titer.

### *E. muscae* is present in the Twyford samples

Having established that DmEV is present in *E. muscae* cells and is replicating at high levels in *D. melanogaster* infected with *E. muscae*, we were curious if we could find evidence of *E. muscae* in samples in which closely related viruses were identified. We began with data from (Webster et al., 2015). We obtained reads for the original samples from the NCBI’s Sequence Read Archive (SRA): SRR1914527 which contains flies from UK (including Twyford) and SRR1914484 which contains flies from non-UK sources.

Using an set of 17,826 genes from a preliminary annotation of the *E. muscae* genome filtered to remove regions that cross-align with the *D. melanogaster* genome, we found 823 read pairs that align to *E. muscae* in the Twyford sample while there are 0 in the non-Twyford sample. An additional 575 read pairs from the Twyford sample align discordantly to the *E. muscae* annotation, while 1,220 have a single read from the pair that aligns, reflecting the incomplete and fragmented nature of the current *E. muscae* annotation. In total reads aligning to 1,500 distinct *E. muscae* genes were identified with an average mismatch frequency of 0.012, consistent with the sample containing a strain of *E. muscae* closely related to but divergent from the Berkeley sample from which the genome was derived.

As described above, (Webster et al., 2015) had noted an unusual profile of small RNAs isolated from the Twyford sample that aligned to the Twyford virus genome. We therefore sequenced small RNAs from our *in vitro* culture. The 924,861 small RNA reads that align to DmEV from our *in vitro E. muscae* culture have very similar properties to those described for Twyford (Figure 4). They show a ~70/30 negative strand bias with a strong preference for a 5’ U base. Furthermore, using Augustus software to predict open reading frames from our de novo-assembled *E. muscae* genome, we see at least one clear Dicer homolog (g4150, e-value < e^-134, % identity > 35%), suggesting that small RNAs may be processed by a Dicer pathway in *E. muscae*.

**Figure 4.**
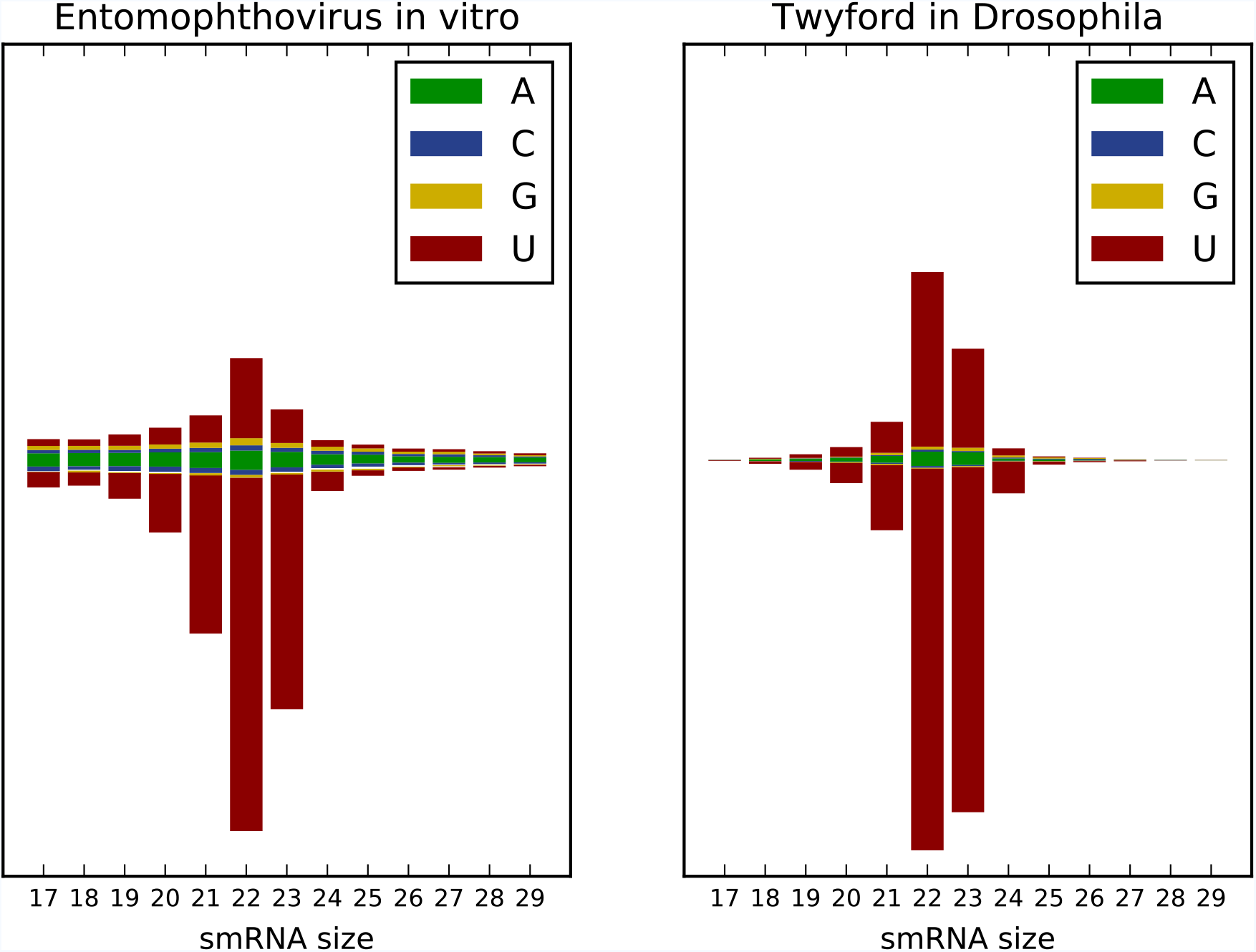
Small RNAs from Entomophthovirus have a characteristic 5’ U bias. We sequenced 204.5 million small RNAs from our *E. muscae in vitro* culture. After quality trimming and adapter removal (using cutadapt (Martin, 2011)), the 17-29 bp fraction was aligned to the entomophtovirus genome (Hisat2 (Kim et al., 2015)). Following the presentation of data from (Webster et al., 2015), the length and 5’ nucleotide bias of aligned reads is shown, with the data from (Webster et al., 2015) for Twyford virus in wild *Drosophila* shown alongside. In both, the aligned reads are mostly 21-23 nt, with a negative strand bias and a strong 5’ U bias.

Collectively this evidence demonstrates that the Twyford virus described by (Webster et al., 2015) as a *D. melanogaster* virus was in fact DmEV present in their sample because of a concurrent infection of at least one of their flies with *E. muscae*.

### *Entomophthovirus* in other samples

GenBank contains a second virus closely related to DmEV and Twyford, Hubei picorna-like virus 39 (H39; KX883974.1). H39 was identified using similar methods to those of (Webster et al., 2015) as part of a large survey of viruses from different collections of arthropod taxa (Shi et al., 2016). We obtained the raw sequencing reads for the 67 different samples used in this experiments and aligned them to both DmEV and the *E. muscae* transcriptome.

The sample from which H39 was identified, a collection of diverse dipterans including one species of *Drosophila*, contains 13,258 reads that align to DmEV as well as 762 reads (out of 96,396,434) that align to 342 different *E. muscae* transcripts. This established that a second wild sample of flies from which a close relative of DmEV was isolated also contained *E. muscae*. None of the remaining 66 samples from (Shi et al., 2016) contain reads that align to either *E. musae* or to any version of DmEV.

Having initially discovered Entomophthovirus (EV) in a shotgun transcriptome assembly, we searched NCBI’s Transcriptome Sequence Assembly (TSA) database for closely related transcripts and identified a series of hits from an individual of *Delia radicum*, a dipteran known as the cabbage fly, infected with *E. muscae* (De Fine Licht et al., 2017). The hits include an essentially full-length EV annotated as an *E. muscae* transcript (GenBank locus GENB01034640), which we henceforth refer to as DrEV. All three wild-caught *E. muscae* infected *D. radicum* had large numbers of reads aligning to DrEV. This dataset also included mRNA sequencing data from six individuals of the housefly *Musca domestica* infected in the laboratory from two wild *M. domestica* infected with *E. muscae*. We identified a different variant of EV in all six of these samples. We refer to this variant as MdEV.

Interestingly, we found no reads aligning to any version of EV in sequencing data from the *in vitro* cultures of *E. muscae* derived from one of the wild caught *M. domestica* described by (De Fine Licht et al., 2017), demonstrating that it is possible to clear the viral infection of *E. muscae*.

Finally, we searched for EV in transcriptome data from flies at various stages of infection with a variety of different pathogenic bacteria to explore whether EV might be an opportunistic infection in flies undergoing immune collapse (Troha et al., 2018) and did not find any, consistent with the observation from (Webster et al., 2015) that Twyford virus is rare in *Drosophlia*.

In summary, our survey of currently published sequencing data suggests that EV is obligately associated with *E. muscae* in *in vivo* infections. We cannot find a single hit for EV and its very close relatives (including Twyford and H39), where *E. muscae* infection was not confirmed phenotypically or suggested by a large number of reads aligning specifically to numerous *E. muscae* transcripts (Shi et al. 2016, Webster et al. 2015). This co-occurrence of fungus and virus appears in a variety of dipteran hosts, including *D. melanogaster*, *M. domestica*, and *D. radicum*.

### Diversity of Entomophthovirus

Detailed analysis of the reads from our *D. melanogaster* samples revealed the presence of three substantially different versions of EV, two (DmEV1 and DmEV2) dominant in the *in vitro* culture, the other (DmEV3) dominant in the *in vivo* samples. DmEV1 and DmEV2 have a pairwise nucleotide divergence of .05, meaning they differ at one base in 20, while these two have an average pairwise divergence of .19 to DmEV3.

Both nucleotide and protein phylogenies (Figure 5) of the seven sequences - DmEV1, DmEV2, DmEV3, Twyford, H39, DrEV and MdEV - place DmEV1 and DmEV2 together with Twyford (which we also assume was derived from *D. melanogaster*) with the three in a clade with DrEV, while DmEV3 is the sister taxa of MdEV in a separate clade. The placement of H39 is inconsistent: the nucleotide tree places H39 as an outgroup to the other six, the protein tree as a deeply branching member of the clade with DmEV1, DmEV2, Twyford and DrEV.

**Figure 5.**
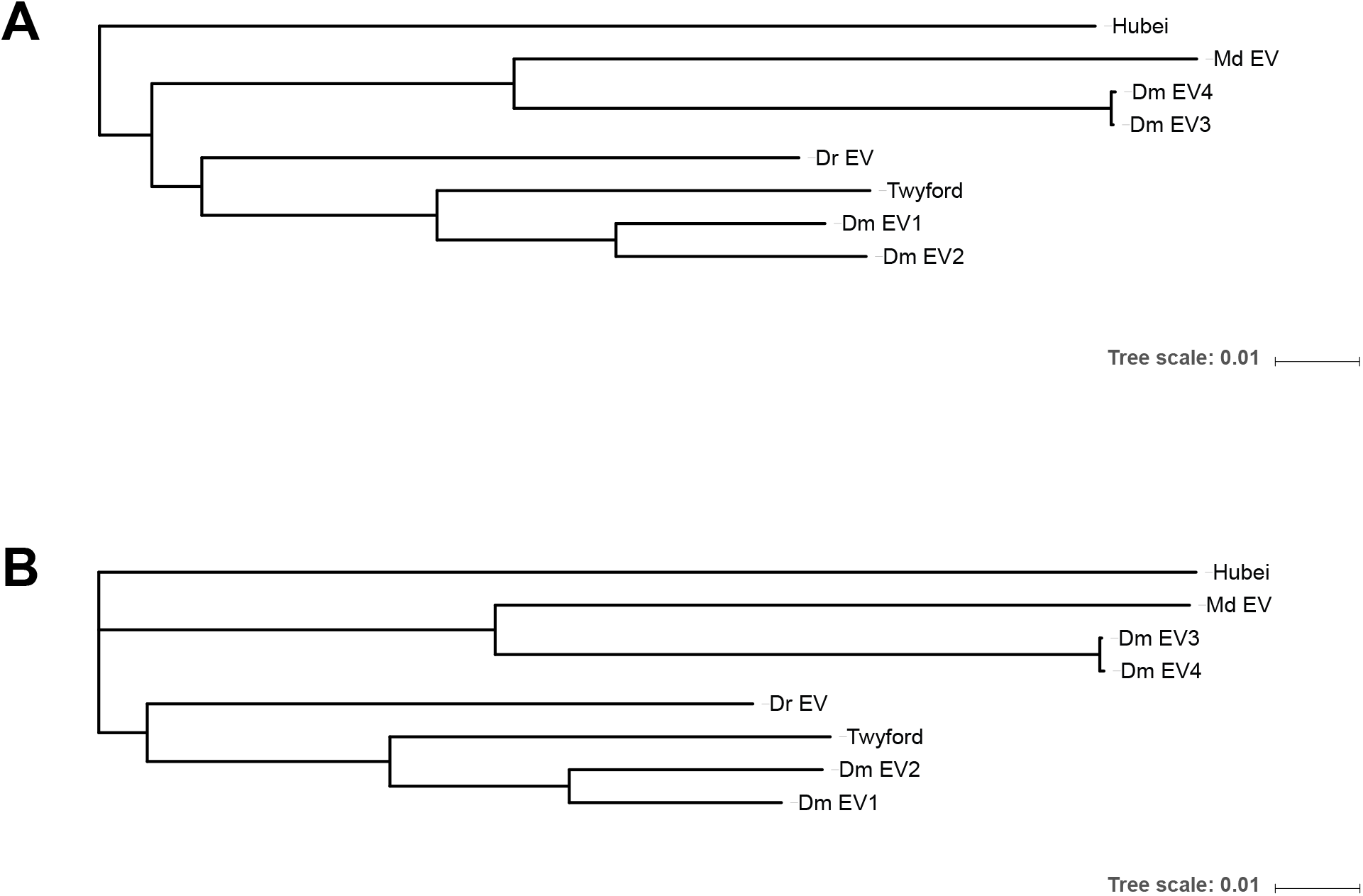
Relationship of *Entomophthora muscae* associated Entomophthovirus variants. Evolutionary relationships of eight Entomophthovirus samples identified from *Entomophthora muscae* infected *Drosophila melanogaster* (Dm and Twyford), *Musca domestica* (Md), *Delia radicum* (Dr) and an unknown dipteran (Hubei) based on (A) nucleotide sequences and (B) inferred amino acid sequences. Trees were calculated using MEGA7 (Kumar et al., 2016) using the Neighbor-Joining method (Saitou and Nei, 1987). The evolutionary distances were computed using the Maximum Composite Likelihood method (Tamura et al., 2004) and are in the units of the number of substitutions per site. The optimal tree is shown, drawn to scale using iTOL (Letunic and Bork, 2016).

Unsurprisingly, we also see evidence for ongoing evolution of the virus. We see a small number of polymorphisms within DmEV3 in our *in vivo* time course. They are too distant from each other to phase, but there are a set of four polymorphisms that are always at the same frequency in the reads from an individual fly but at different frequencies between individuals, suggesting there is some form of bottlenecking of a mixed population, likely during spore transmission or the early phases of infection, although we have not demonstrated this experimentally.

## Discussion

We believe these data unambiguously establish that the virus we originally identified in our *E. muscae in vitro* culture, along with two closely related viruses previously described as dipteran viruses, are variants of an iflavirus that infects *E. muscae* and is transmitted along with *E. muscae* to and from infected dipteran hosts. We have demonstrated that the virus is actively replicating in *E. muscae* in culture, that *E. muscae* generates anti-viral RNAs from the virus, that the virus forms capsids in the fungus, that it is transmitted along with the fungus from fly to fly, and that where the virus is found in nature it is always associated with *E. muscae* infected dipterans. For these reasons we formally propose naming this virus Entomophthovirus.

This is, to our knowledge, the first known member of the viral order Picornavirales to infect a fungus. As virtually all other known iflaviruses are insect pathogens, it seems likely that the virus moved from an insect host to an ancestor of *E. muscae*. When this happened and how broad the association is remains to be seen, but there is precedent for host switching in iflaviruses (Saqib et al., 2015), and many fungal viruses are members of families that also infect animals (Son et al., 2015).

The consistent association suggests either that, in spite of an active anti-viral response from the fungus, the virus is never cleared from fungal cells and is present in fungal spores. The extent of virus-associated fungal mortality is unclear, but the high levels of viral RNA and the large number of capsids in the *in vitro* culture suggest that the virus has relatively low pathogenicity.

An alternative hypothesis is that EV provides some fitness advantage to *E. muscae* and its presence is essential to successful transmission of the infection, serving as a form of positive selection to maintain the association. One tantalizing possibility is that the virus is involved in behavior manipulation. Many animal viruses have behavioral effects on their hosts, including a baculovirus that induces summiting behavior in caterpillars (Katsuma et al., 2012), and, more specifically several other iflaviruses that induce a range of behaviors in their insect hosts (Dheilly et al., 2015; Fujiyuki et al., 2005).

For example, Kakugo Virus, which is a subtype of Deformed-Wing Virus, is an iflavirus that infects honeybees, and has been shown to be associated with aggressive colony behavior (Fujiyuki et al., 2005). Another iflavirus (*D. coccinellae* paralysis virus; DcPV) was recently shown to be actively involved in the parasitoid induced paralytic behavioral manipulation of ladybugs (Dheilly et al., 2015). Finally, an iflavirus was recently found associated with *Bombyx mori* infected with the behavioral manipulating ascomycete fungus *Cordyceps militaris (Suzuki et al., 2015)*, although the nature of the fungal-viral association and its significant are unknown.

Many important questions regarding the relationship between EV and *E. muscae* remain open. Can *E. muscae* cleared of virus infect flies, induce behaviors and transmit the infection between flies? Does the virus replicate in fly cells during infection, or is it restricted to the fungus? Can the virus infect flies in the absence of fungus, and, if so, does it induce behavioral changes? How widespread is the association between entomopathogenic fungi and viruses?

On this later point, another distantly related picorna-like virus, RiPV-1, was recently identified in mRNA sequencing reads from bean bugs, *Riptortus pedestris*, infected with a distantly related ascomycete gentomopathogenic fungus, *Beauveria bassiana* (Yang et al., 2016). Like EV, RiPV-1 replicates at high rate in infected insects. In this case the authors concluded that the virus is not restricted to fungal infected animals, rather that there is a persistent low-level infection in their laboratory stocks. Nonetheless, the observation is intriguing and may suggest a broader relationship between fungal insect pathogens and positive-strand RNA viruses.

Whether it turns out the EV is involved in behavioral manipulation or not, it is a fascinating example of viral adaptation that represents the discovery of the expansion of a major viral lineage to a new kingdom, and understanding how this relationship evolved and is maintained will illuminate new aspects of virus biology.

## Methods

### Confirming virus in samples by RT-PCR

A liquid culture of *E. muscae* was propagated in Grace’s Insect Media supplemented with 5% FBS. One milliliter of cultured cells was spun at 10,000 x g for 5 min to pellet fungal cells. The supernatant was filtered through a 0.22 um syringe filter, and RNA was extracted with Trizol. The pellet was resuspended in 10 mL PBS and vacuum filtered through a Whatman 0.8 um cellulose ester filter to collect cells. This filter was washed four times with 10 mL of PBS. RNA was extracted from all washes with Trizol. Finally the filter paper was equilibrated in 10 mL PBS for 30 minutes and the eluted cells were pelleted and RNA-extracted by Trizol. Additionally, Trizol extraction was performed on our media stocks and 25 CantonS flies from the fly stocks we have used to propagate *in vivo E. muscae* infections.

All RNA samples were reverse transcribed with SuperScript III reverse transcriptase using 150 ng of random hexamer primers per reaction. The RT reaction was heat inactivated at 70C for 15 minutes and 1/10 of the cDNA was used to amplify a 831-bp sequence specific to EV using Taq polymerase. Amplification primers were “GGGTTAGAAGTGTGCGAGAAT” and “GCGACAAGGACTACACGATAAG”. Amplicon presence was assayed with a 1% agarose gel with ethidium bromide.

### Analysis of *E. muscae* infected *D. melanogaster* RNA-seq

We used RNA-seq data from (Elya et al., 2017) available in the NCBI GEO database under ID GSE111046. We used published read counts for each sample from (Elya et al., 2017) for *E. muscae* and *D. melanogaster*, and determined read counts for EV by aligning reads to all variants of the EM genome using bowtie2 (Langmead, 2010).

### Transmission Electron Microscopy

To prepare a crude sample of extracellular EV, we pelleted 10 mL of *E. muscae* liquid culture, filtered the supernatant through a 0.2 um syringe filter, and ultra-centrifuged the sample for 2 hours at 25,000 RPM and 4C, using the SW28 swinging bucket rotor. The pelleted material was fixed in 2.5% glutaraldehyde (in 0.1M sodium cacodylate, pH 7.4) and a 1:100 dilution was negative stained with 1% uranyl acetate and imaged on a Tecnai 12 TEM.

To image sections of *E. muscae* cells, pelleted cells were fixed in 2.5% glutaraldehyde in 0.1M sodium cacodylate, pH 7.4 and then embedded in 2% agarose. After washing away fixation buffer with 0.1M sodium cacodylate, samples were treated with 1% osmium tetroxide and 1.6% potassium ferricyanide for 30 minutes, then washed again with 0.1M sodium cacodylate. Fixed and embedded samples were then dehydrated with increasing concentrations of EtOH, followed by pure EtOH and then pure acetone. Increasing concentrations of Eponate resin in acetone (25%, 50%, 75%, then 100%) were infiltrated into the samples for one hour each, followed by pure resin infiltration overnight.. Then eponate resin with BDMA accelerant was infiltrated into samples for 5 hours. Samples were embedded into a mold and incubated ta 60C for one week. 70 nm sections of samples were cut and stained with 2% uranyl acetate, followed by Reynolds lead citrate, before imaging on the Tecnai 12 TEM.

### Small RNA sequencing

4 mL of *E. muscae* liquid culture was pelleted and RNA was extracted with Trizol. The sample was treated with Turbo DNase and Trizol-extracted again. RNA integrity was confirmed with an RNA 6000 Pico chip on the Agilent 2100 Bioanalyzer. RNA was diluted to 200 ng/ul, and 1 ug (5 ul) was used as input for the Illumina TruSeq small RNA kit (RS-200-0012). The size range was confirmed on the 2100 Bioanalyzer with a HS DNA chip. 204.5 million 50 SR reads were obtained with a HiSeq 4000.

Cutadapt software was used to trim 3’ bases with a Phred score <= 10, and to remove the 3’ Illumina small RNA adapter. Next, cutadapt was used to select sequences between 17 and 29 bp and these reads were aligned to the EV genome with Hisat2.

### Analysis of samples with previously identified Entomophthovirus

#### Twyford

We obtained reads for the original samples from the NCBI’s Sequence Read Archive (SRA): SRR1914527 which contains flies from UK (including Twyford) and SRR1914484 which contains flies from non-UK sources. We aligned reads using bowtie2 with default parameters to a set of 17,826 genes from a preliminary annotation of the *E. muscae* genome filtered to remove regions that cross-align with the *D. melanogaster* genome (identified using blastn with an e-value cutoff of .0000001) and highly conserved fungal genes which have regions that align cross species (e.g. beta-tubulin, histones, ribosomal proteins), we found 823 read pairs that align to *E. muscae* in the Twyford sample while there are 0 in the non-Twyford sample. An additional 575 read pairs from the Twyford sample align discordantly to the *E. muscae* annotation, while 1,220 have a single read from the pair that aligns, reflecting the incomplete and fragmented nature of the current *E. muscae* annotation. In total reads aligning to 1,511 distinct *E. muscae* genes were identified with an average mismatch frequency of .012. Six reads align from SRR1914484 align to the filtered *E. muscae* transcriptome, but have a high number of mismatches demonstrating they are not from *E. muscae*. 807 reads aligned to EV from SRR1914527 with an average mismatch frequency of .0036. Four reads from SRR1914484 aligned to the highly conserved RDRP portion of EV with mismatch patterns suggesting they are from a different virus.

#### H39

We obtained reads from all 67 samples from the experiment in which H39 was identified (Shi et al., 2016) (NCBI SRA BioProject ID SRP073469) and aligned with the same procedure as for Twyford above. H39 was identified in sample SRR3400838 labeled “Diptera mix”. It contains 13,258 reads that align to EV and 762 reads that align to 342 different *E. muscae* genes. None of the remaining 66 samples appear to contain *E. muscae*. 15 have a handful of reads (between 1 and 18) that align to the filtered *E. muscae* transcript set, but in all cases they contain many mismatches (from 10 to 25 per 100bp) demonstrating that the reads are from another fungal species. Four samples have a small number of reads (1-25) aligning to the conserved RDRP, but with multiple mismatches showing that they are no EV. To confirm that these samples do not contain EV we assembled the reads from these samples using TRINITY and did not find any even fragmentary versions of EV in the assemblies.

### Identification of *Delia radicum* and *Musca domestica* Entomophthovirus

We searched NCBI’s Transcriptome Sequence Assembly (TSA) database for additional and identified a series of hits from an individual of *Delia radicum*, a dipteran known as the cabbage fly, infected with *E. muscae* (De Fine Licht et al., 2017). The hits include an essentially full-length EV annotated as an *E. muscae* transcript (GenBank locus GENB01034640), which we henceforth refer to as DrEV.

We downloaded reads for this paper (NCBI SRA BioProject ID PRJEB10825), which involved the sampling of two infected individuals of the house fly *Musca domestica* and three of *D. radicum*. Spores from the first *M. domestica* sample were used to infect *M. domestica* in the laboratory, and mRNA from three laboratory-infected *M. domestica* were sequenced (A_Md1, A_Md2, and A_Md3) and a fourth was used to inoculate three *in vitro* cultures, from which RNA was also sequenced (yielding mRNA samples A_Gl1, A_Gl2, and A_Gl3). The first part of the process was repeated for a second wild infected *M. domestica* (yielding mRNA samples B_Md1, B_Md2 and B_Md3). Finally mRNA from three wild caught individuals of *D. radicum* were sequenced (yielding mRNA samples C_Dr, D_Dr, E_Dr).

All three wild *E. muscae* infected *D. radicum* samples contain EV. One wild *D. radicum* had 50,000 EV reads, while the other two had only a few hundred. Preliminary alignments of the *M. domestica* samples to *D. melanogaster* EV showed a wide range of EV titres: laboratory *M. domestica* infected from the first individual had only approximately 100 EV reads each, while laboratory *M. domestica* infected from the second wild individual had several hundred thousand EV reads.

We used reads from *M. domestica* infected by the second wild *M. domestica* sample to assemble to *de novo* transcriptome using Trinity (Grabherr et al., 2011). This transcriptome contains a nearly full length copy of EV which we refer to as MdEV. We aligned the *M. domestica* samples against MdEV confirming the presence of this virus in all six infected *M. domestica*, and its absence from the *in vitro* culture.

## Acknowledgements

We are grateful to Joseph DeRisi for helpful discussions about the identification, isolation and characterization of viruses. We thank Danielle Jorgens and Reena Zalpuri at the UC Berkeley Electron Microscopy Lab. This work was funded by an HHMI Investigator award to MBE. CNE was supported by an NSF predoctoral fellowship.

## References

Balazy, S. (1984). On Rhizoids of Entomophthora-muscae (Cohn) Fresenius (Entomophthorales, Entomophthoraceae). Mycotaxon 19, 397–407.

Cohn, F. (1855). Empusa muscae und die krankheit der Stubenfliegen. Ein Beitrag zur Lehre von den durch parasitische Pilze charakterisierten Epidemien. Nova Acta Academiae Caesareae Leopoldino-Carolinae Germanicae Naturae Curiosorum 25, 299–360.

De Fine Licht, H.H., Jensen, A.B., and Eilenberg, J. (2017). Comparative transcriptomics reveal host-specific nucleotide variation in entomophthoralean fungi. Mol. Ecol. 26, 2092–2110.

Dheilly, N.M., Maure, F., Ravallec, M., Galinier, R., Doyon, J., Duval, D., Leger, L., Volkoff, A.-N., Missé, D., Nidelet, S., et al. (2015). Who is the puppet master? Replication of a parasitic wasp-associated virus correlates with host behaviour manipulation. Proceedings of the Royal Society of London B: Biological Sciences 282, 20142773.

Eisthen, H.L., and Theis, K.R. (2016). Animal-microbe interactions and the evolution of nervous systems. Philos. Trans. R. Soc. Lond. B Biol. Sci. 371, 20150052.

Elya, C., Lok, T.C., Spencer, Q.E., McCausland, H., and Eisen, M.B. (2017). A fungal pathogen that robustly manipulates the behavior of Drosophila melanogaster in the laboratory.

Forsythe, P., Kunze, W.A., and Bienenstock, J. (2012). On communication between gut microbes and the brain. Curr. Opin. Gastroenterol. 28, 557–562.

Fujiyuki, T., Takeuchi, H., Ono, M., Ohka, S., Sasaki, T., Nomoto, A., and Kubo, T. (2005). Kakugo virus from brains of aggressive worker honeybees. Adv. Virus Res. 65, 1–27.

Grabherr, M.G., Haas, B.J., Yassour, M., Levin, J.Z., Thompson, D.A., Amit, I., Adiconis, X., Fan, L., Raychowdhury, R., Zeng, Q., et al. (2011). Full-length transcriptome assembly from RNA-Seq data without a reference genome. Nat. Biotechnol. 29, 644–652.

Hajek, A.E., Papierok, B., and Eilenberg, J. (2012). Chapter IX - Methods for study of the Entomophthorales. In Manual of Techniques in Invertebrate Pathology (Second Edition), L.A. Lacey, ed. (San Diego: Academic Press), pp. 285–316.

Hoover, K., Grove, M., Gardner, M., Hughes, D.P., McNeil, J., and Slavicek, J. (2011). A Gene for an Extended Phenotype. Science 333, 1401–1401.

Hughes, D.P., Araújo, J.P.M., Loreto, R.G., Quevillon, L., de Bekker, C., and Evans, H.C. (2016). From So Simple a Beginning: The Evolution of Behavioral Manipulation by Fungi. Adv. Genet. 94, 437–469.

Katsuma, S., Koyano, Y., Kang, W., Kokusho, R., Kamita, S.G., and Shimada, T. (2012). The baculovirus uses a captured host phosphatase to induce enhanced locomotory activity in host caterpillars. PLoS Pathog. 8, e1002644.

Kim, D., Langmead, B., and Salzberg, S.L. (2015). HISAT: a fast spliced aligner with low memory requirements. Nat. Methods 12, 357–360.

Kramer, J.P., and Steinkraus, D.C. (1981). Culture of Entomophthora muscae in vivo and its infectivity for six species of muscoid flies. Mycopathologia 76, 139–143.

Kumar, S., Stecher, G., and Tamura, K. (2016). MEGA7: Molecular Evolutionary Genetics Analysis Version 7.0 for Bigger Datasets. Mol. Biol. Evol. 33, 1870–1874.

Langmead, B. (2010). Aligning short sequencing reads with Bowtie. Curr. Protoc. Bioinformatics Chapter 11, Unit 11 7.

Letunic, I., and Bork, P. (2016). Interactive tree of life (iTOL) v3: an online tool for the display and annotation of phylogenetic and other trees. Nucleic Acids Res. 44, W242–W245.

Libersat, F. (2003). Wasp uses venom cocktail to manipulate the behavior of its cockroach prey. J. Comp. Physiol. A Neuroethol. Sens. Neural Behav. Physiol. 189, 497–508.

Martin, M. (2011). Cutadapt removes adapter sequences from high-throughput sequencing reads. EMBnet.journal 17, 10–12.

Rohrscheib, C.E., and Brownlie, J.C. (2013). Microorganisms that Manipulate Complex Animal Behaviours by Affecting the Host’s Nervous System. Springer Science Reviews 1, 133–140.

Roy, H.E., Steinkraus, D.C., Eilenberg, J., Hajek, A.E., and Pell, J.K. (2006). Bizarre interactions and endgames: entomopathogenic fungi and their arthropod hosts. Annu. Rev. Entomol. 51, 331–357.

Saitou, N., and Nei, M. (1987). The neighbor-joining method: a new method for reconstructing phylogenetic trees. Mol. Biol. Evol. 4, 406–425.

Sampson, T.R., and Mazmanian, S.K. (2015). Control of Brain Development, Function, and Behavior by the Microbiome. Cell Host Microbe 17, 565–576.

Saqib, M., Wylie, S.J., and Jones, M.G.K. (2015). Serendipitous identification of a new Iflavirus-like virus infecting tomato and its subsequent characterization. Plant Pathol. 64, 519–527.

Shi, M., Lin, X.-D., Tian, J.-H., Chen, L.-J., Chen, X., Li, C.-X., Qin, X.-C., Li, J., Cao, J.-P., Eden, J.-S., et al. (2016). Redefining the invertebrate RNA virosphere. Nature.

Silva, L.A., Ardisson-Araujo, D.M.P., Tinoco, R.S., Fernandes, O.A., Melo, F.L., and Ribeiro, B.M. (2015). Complete genome sequence and structural characterization of a novel iflavirus isolated from Opsiphanes invirae (Lepidoptera: Nymphalidae). J. Invertebr. Pathol. 130, 136–140.

Son, M., Yu, J., and Kim, K.-H. (2015). Five Questions about Mycoviruses. PLoS Pathog. 11, e1005172.

Steinkraus, D.C., and Kramer, J.P. (1987). Susceptibility of sixteen species of Diptera to the fungal pathogen Entomophthora muscae (Zygomycetes: Entomophthoraceae). Mycopathologia 100, 55–63.

Suzuki, T., Takeshima, Y., Mikamoto, T., Saeki, J.-D., Kato, T., Park, E.Y., Kawagishi, H., and Dohra, H. (2015). Genome Sequence of a Novel Iflavirus from mRNA Sequencing of the Pupa of Bombyx mori Inoculated with Cordyceps militaris. Genome Announc. 3.

Tamura, K., Nei, M., and Kumar, S. (2004). Prospects for inferring very large phylogenies by using the neighbor-joining method. Proc. Natl. Acad. Sci. U. S. A. 101, 11030–11035.

Troha, K., Im, J.H., Revah, J., Lazzaro, B.P., and Buchon, N. (2018). Comparative transcriptomics reveals CrebA as a novel regulator of infection tolerance in D. melanogaster. PLoS Pathog. 14, e1006847.

Wang, G., Zhang, J., Shen, Y., Zheng, Q., Feng, M., Xiang, X., and Wu, X. (2015). Transcriptome analysis of the brain of the silkworm Bombyx mori infected with Bombyx mori nucleopolyhedrovirus: A new insight into the molecular mechanism of enhanced locomotor activity induced by viral infection. J. Invertebr. Pathol. 128, 37–43.

Webster, C.L., Waldron, F.M., Robertson, S., Crowson, D., Ferrari, G., Quintana, J.F., Brouqui, J.-M., Bayne, E.H., Longdon, B., Buck, A.H., et al. (2015). The Discovery, Distribution, and Evolution of Viruses Associated with Drosophila melanogaster. PLoS Biol. 13, e1002210.

Yang, Y.-T., Nai, Y.-S., Lee, S.J., Lee, M.R., Kim, S., and Kim, J.S. (2016). A novel picorna-like virus, Riptortus pedestris virus-1 (RiPV-1), found in the bean bug, R. pedestris, after fungal infection. J. Invertebr. Pathol. 141, 57–65.

Zuckerkandl, E., and Pauling, L. (1965). Evolutionary Divergence and Convergence in Proteins. In Evolving Genes and Proteins, V. Bryson, and H.J. Vogel, eds. (Academic Press), pp. 97–166.

